# Category-specific fMRI correlates of picture naming: A study with Arabs and Filipinos

**DOI:** 10.1101/2022.02.05.478294

**Authors:** Haythum O. Tayeb, Jamaan Alghamdi, Naushad Ahmed, Yousef Alsawwaf, Khalid Alsafi, Abrar Baduwailan, Bassam Yaghmoor, Tariq Elyas, Mohammed Mudarris, Daniel S. Weisholtz

## Abstract

Cross cultural neuroimaging work has demonstrated differences in neural correlates of some cognitive processes between individuals from different cultures, often comparing American and Chinese subjects. In contrast, a limited number of studies examined Arab and/or Filipino participants. This fMRI study aimed to demonstrate neural activations during animal and tool picture naming by 18 healthy Arabs and 18 healthy Filipino participants. In animal naming contrasted with tool naming, Arabs preferentially activated regions in the right lateral occipital and fusiform cortices, whereas Filipinos recruited bilateral visual areas. Cross-group comparisons of animal naming revealed that Arabs recruited right visual areas more than Filipinos, who in turn recruited the cerebellum more than Arabs. In tool naming, Arabs preferentially activated a predominantly left frontoparietal network, whereas no regions were identified in Filipinos, and no differences in activation between groups were found. Using a low-demand picture-naming task, this study revealed category-specific neural activations during picture naming by Arabs and Filipinos, as well as between-group differences in animal naming. The results suggest that Arabs and Filipinos may have culture-specific differences in processing animate and inanimate pictures, and caution against generalizing findings from the more commonly studied populations, especially in verbal tasks such as picture naming.

**HIGHLIGHTS:**

▪ The neural correlates of animal and tool picture naming in Arabs and Filipinos are category specific.
▪ Animal naming by Arabs tended to preferentially activate the nondominant ventral visual stream.
▪ Animal naming by Filipinos activated bilateral visual areas, and the cerebellum.
▪ Tool naming by Arabs activated dominant frontoparietal areas related to praxis.
▪ Results suggest that Arabs and Filipinos have culture-specific differences in visual processing.

## INTRODUCTION

Neural correlates of semantic processing of visual stimuli may vary across dissimilar cultures (Han et al., 2013; Kovelman et al., 2008; Nisbett & Miyamoto, 2005; Storbeck et al., 2006). Prior work compared East Asian and Western subjects during visual perceptual tasks with regards to their attentional strategies and their associated neural mechanisms while processing visual scenes. American participants focused on details of objects, whereas Chinese participants devoted more attention to contexts and relationships (Jenkins et al., 2010; Kuwabara & Smith, 2012). Such mechanisms have not been demonstrated in Arab subjects. A small number of fMRI studies with Arab participants exist (Bourisly et al., 2013; Mohtasib et al., 2021; Sakr, 2011; Sakr & Ali, 2011) describing fMRI use in planning neurosurgical procedures (Sakr, 2011; Sakr & Ali, 2011). Similarly, the limited number of fMRI studies on Filipino participants (or Tagalog speakers) included a number Filipino participants as part of a cross cultural examination, but did not focus on this group (Nauchi & Sakai, 2009; Plante et al., 2015). However, foundational research mapping the neural correlates of cognitive tasks in Arabs and comparing them with other cultures has not been reported.

Arabic, as with other Semitic languages, differs in the morphological complexity from Indo-European languages, such as English (Perea et al., 2014). Arabic morphology, like Hebrew, involves derivations of triconsonantal roots, that are inherently complex as words consist of a combination of morphemes and phonological patterns (Abu-Rabia, 2001). Whereas, English involves prefixes and suffixes, which constitute a predictable derivations from a common morpheme (Bick et al., 2011; Boudelaa & Marslen-Wilson, 2001; Palti et al., 2007). Neuroimaging studies investigating morphological processing in Hebrew reveal a network that is separate from semantic processing, indicating an underlying basis for word meaning (Bick et al., 2010, 2011). This does not seem to be the case in studies examining English language, as morphemes were not clearly distinguishable from semantic or orthographic roots (Bick et al., 2011).

Confrontational picture naming has been shown to differentially activate neural pathways depending on the semantic category of the pictures shown (Chouinard & Goodale, 2010). The most consistent finding in this area is the difference in neural activation between animate and inanimate objects. Naming animate objects (e.g., animals) is associated with greater activations in areas predominantly located in the ventral visual pathway, including the bilateral lateral occipital cortices (OC), right middle occipital gyrus, the posterior fusiform cortex (FFC) bilaterally. In addition, naming animals was associated with greater activations in the left anterior cingulate cortex (ACC), and left cuneus. Naming inanimate objects (e.g., tools), is associated with greater activations in sensorimotor and praxis-related frontoparietal regions. Patients with neurological lesions may manifest category-specific impairments in naming (Capitani et al., 2003). Arab patients with such category-specific impairments have not been reported in the literature, nor have the neural correlates of category-specific semantic processing in Arab subjects. Likewise, studies examining the neural correlates of picture naming in Tagalog speaking Filipino subjects are lacking, with some studies examining picture naming focusing on bilingualism in Tagalog-English speakers (Gollan & Acenas, 2004), and neuroimaging studies examining linguistic properties unique to Tagalog (Wray et al., 2019).

As most cross-cultural studies in this domain have been conducted between American and Chinese speakers, this study aimed to explore neural correlates of picture naming in Arabic speakers, an understudied language with verb-tenses as in the English language, compared to a local population of Tagalog speakers, a tense-less language, similar to Chinese (Dery, 2009). Studies comparing Chinese and American participants on a perceptual picture task, revealed distinct neural-correlates between the groups that were attributed to culture-specific experiences. Specifically that Chinese participants showed greater activation in frontal regions attributed to Asian-specific focus on object-relations, whereas greater temporal and cingulate activation in American subjects was attributed to the American culture’s emphasis on individuals (Gutchess et al., 2006). Such findings put into question whether neural correlates of picture naming may be generalizable based on studies on more frequently studied populations, as cultural-lingual properties may reveal varying neural processing during cognitive and perceptual tasks. This is especially true in tasks with a prominent verbal component, as in the case of confrontational naming.

The main aim of this fMRI study is to demonstrate cerebral activations in healthy Arabs during a confrontational naming task using pictures of animals and tools. In addition, the study will explore differences in activations between healthy Arabs and healthy Filipino subjects doing the same task, based on previously reported differences in brain activity across cultures, and languages for confrontational naming tasks. The fMRI protocol will employ an alternating block-design to investigate picture naming, as such design is suited to detect activity in regions of interest (ROI) in relation to tasks, and have higher statistical power compared to event-related designs, due to better signal-to-noise ratio (Petersen & Dubis, 2012). Event related designs are preferred in aphasic patient studies to be able to separate incorrect trials, using an overt naming task to be able detect errors (Wilson et al., 2017). However, as this study aims to examine activity in healthy participants, a covert (silent) naming task is used to reduce potential artifacts, such as orofacial musculature movement (Chouinard & Goodale, 2010).

## MATERIALS AND METHODS

### Participants

The study participants consist of 18 Arabs (9 males; *M* = 37.56; *SD* = 10.85) and 18 Filipinos (9 males; *M* = 42.18; *SD* = 7.66), all right-handed. The mean age for both cohorts 39.8 (*SD* = 9.59). All participants had at minimum Bachelor level education (16 years), and Master level education at the highest level (18 years). There were no significant differences between Arab and Filipino participants with regards to age or years of education (*p* > .05). The recruited participants were healthy university hospital employees who met the following inclusion criteria: i) All subjects lived in Jeddah, Saudi Arabia at the time of the study and worked at King Abdulaziz University Hospital, a tertiary care academic hospital; ii) Their ages ranged between 20 and 60 years; iii) The Filipino cohort grew up and received education in the Philippines, spoke fluent Tagalog as their first language, and have not lived in Arab countries for longer than 3 years; iv) The Arab cohort grew up and received education in Saudi Arabia, spoke fluent Arabic as their first language, were not fluent in another language, and have not lived outside Saudi Arabia for more than 3 years; iv) All subjects had no medical, neurological, visual or psychiatric problems, took no medications, and had no contraindications to fMRI scanning; v) Subjects agreed to participate in the study and signed a written informed consent form. Subjects from the two groups were matched for gender, age, and education level. The study was approved by the Institutional Review Board at the Faculty of Medicine at King Abdulaziz University, Jeddah, Saudi Arabia.

### Stimuli

The stimuli were selected from a picture bank that has been extensively examined in research of neural correlates of picture naming tasks (Snodgrass & Vanderwart, 1980). Forty animal, and forty tool pictures were presented to the participants in a randomized block design. The pictures were of a similar size, with a resolution of 960×720 pixels, which consisted of black line drawings presented in the center of a white background, subtending a central visual angle of 20 degrees. The lexical frequency of the words have been reported for both Arabic (Boudelaa & Marslen-Wilson, 2010), and Tagalog (Baisa et al., 2019; Kilgarriff et al., 2014; Zuraw, 2010; Supplementary Table S1, and S2). A list of the stimuli names, and lexical frequencies are listed in the supplementary materials (Table S1, and S2). Lexical frequencies for names of the Arabic animal pictures (*M =* 4.07, *SD =* 10.81, Range = 0.03 – 64.22) were significantly lower *(t(38)* = -2.84, *p* < 0.01, Cohen’s *d* = -0.66) than the lexical frequency of the Tagalog names for the same stimuli (*M =* 21.66, *SD =* 36.18, Range = 0.04 – 174.26). Similarly, the lexical frequencies of the names for the Tool picture names (*M =* 5.86, *SD =* 13,82, Range = 0.03 – 78.60) were significantly lower (*t(38)* = -2.27, *p* = .03, Cohen’s *d* = - 0.54) than those reported for Tagalog (*M =* 16.26, *SD =* 23.79, Range = 0.05 – 83.64). While subjective psycholinguistics properties (e.g., name agreement, familiarity) have been reported for Tunisian and Qatari Arabic speaking participants (Boukadi et al., 2016; Khwaileh et al., 2018), to the best of our knowledge, similar findings have not been reported for Tagalog speakers.

### fMRI and Experimental Paradigms

Using a 3-tesla Siemens MAGNETOM Skyra MRI scanner (TE=30 ms, TR=3000 ms, flip angle=90°, field of view=94 × 94, number of slices=36, slice thickness=3mm), we obtained fMRI scans while subjects viewed and named images of either animals or tools. Paddings were placed on either side of the participant’s head to reduce movement by the participant to a minimum. Each scanning session was divided into two runs, each run containing nine blocks, with each block lasting 30 seconds. During the first run, 5 blocks were rest blocks, during which subjects stared at a crosshair display. These alternated with four testing blocks, during each of which subjects silently named 10 successively-appearing pictures of animals (figure 1). Each picture was shown for 3 seconds. The fMRI technician communicated with the participant between each run to ensure that the participant remained awake and comfortable throughout the testing procedure. The second run was identical to the first run except that the pictures shown during the testing blocks were of tools. The chosen pictures were commonly used for confrontation naming, and have been used in similar studies (Snodgrass & Vanderwart, 1980).

**Figure (1):**
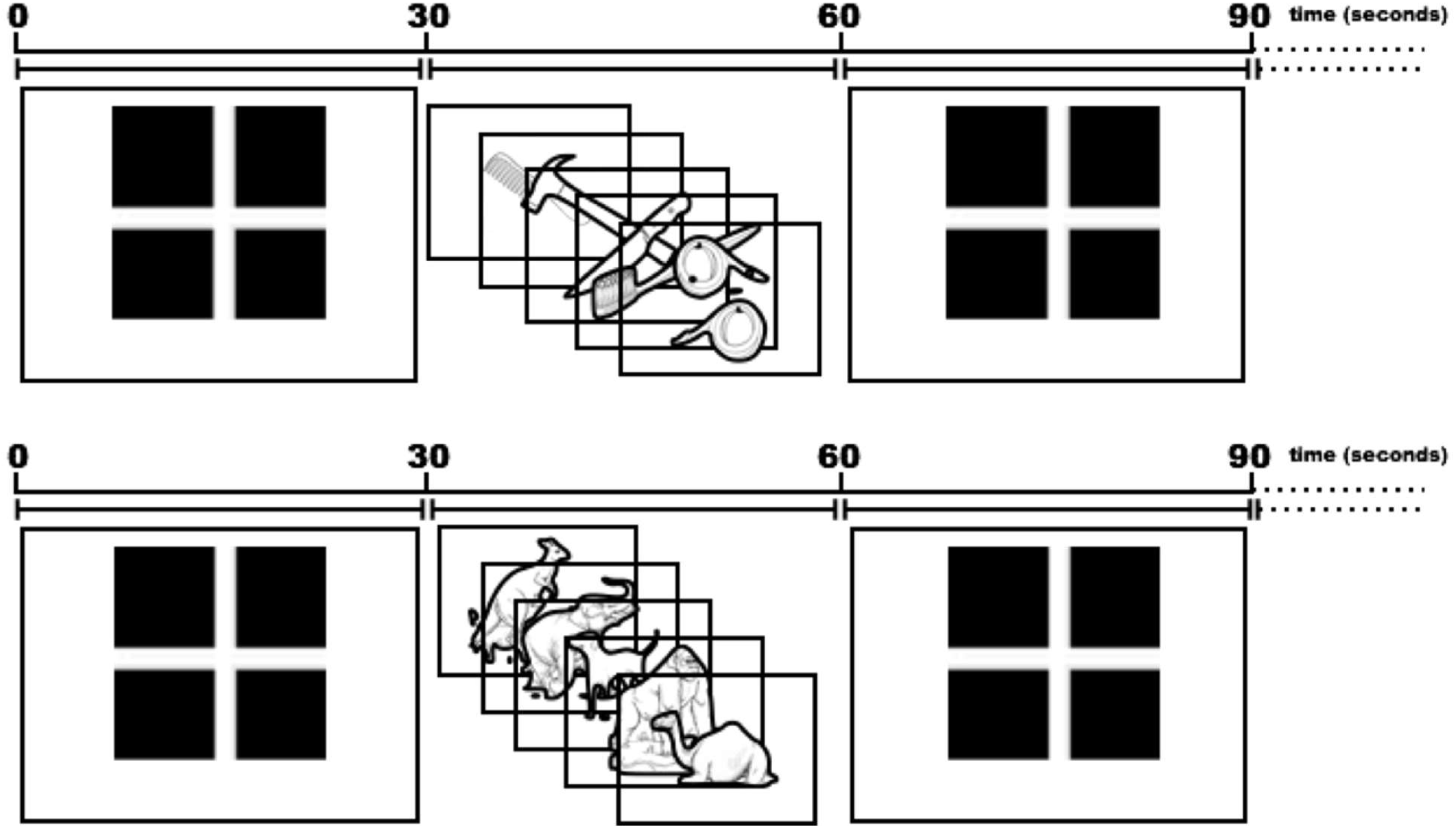
Schematic representation of the fMRI paradigm used in the study

The collected scans were preprocessed and analyzed using FMRIB Software Library (FSL) software. Functional data were preprocessed to remove low frequency linear drift in the signal from each functional data set (Jenkinson et al., 2012). Motion correction, temporal filtering and spatial smoothing (5mm FWHM Gaussian window) of the data were also applied. Then, registration to a T1 space image; and (5) FSL/FNIRT-based registration to MNI 2-mm space were conducted. For this aspect of data preprocessing, standard filtering techniques available in FSL were applied. Individual activation maps from each participant were obtained. The size of the activated area was calculated from the volume of activation exceeding a significance threshold of *p* < .001 and forming a cluster significance of *p* < .05.

Paired t-test was applied to showcase the distinct areas of activation associated with each task. Task activations were compared within group in contrast to baseline activation, and then across groups. Finally, cluster-thresholding was carried out to reveal clusters that were significantly activated, with a *p*-value significance threshold set to .05. All regions were identified using the Harvard-Oxford cortical and subcortical structural atlases (within FSL).

The lateralization of brain activity has been evaluated by calculating the laterality index (LI) for each participant. It was calculated as the proportion of active voxels in the left versus the right region of interest (ROI) averaged across multiple thresholds (Ito & Liew, 2016; Jansen et al., 2006; Seghier, 2008). We set a threshold with range of z-values (z = 1.0, z = 1.5, z = 2.3) to test the effects of different thresholds on the second-level whole brain map using FSL cluster tool. Then we used fslstats tool to determine the total number of active voxels in all ROIs in both hemispheres. Finally, we calculated LI using the formula: LI = (left − right) / (left + right). This yields a score for LI range from +1 (all left hemisphere activation only) to −1 (all right hemisphere activation only) and the intermediate values reflect varying degrees of laterality (Jansen, Menke et al., 2006).

## RESULTS

Cerebral regions showing statistically-significant blood oxygen-level dependent (BOLD) signal increases during each of the animal and tool picture naming tasks in comparison to the rest condition in Arab and Filipino participants are shown in Tables 1 and 2, and Figures 2 and 3. Cross-task comparisons are shown in Table 3 and Figure 4. During animal naming, Arabs more significantly recruited the right lingual gyrus, the right fusiform cortex (FFC), along with the bilateral occipital poles (OPs) in comparison to when they named tools. During tool naming, Arabs more significantly recruited the bilateral superior parietal lobules (SPLs), and the left postcentral gyrus in comparison to when they named animals. Filipinos appeared to activate bilateral visual areas during animal naming in comparison to tool naming. Cross-group comparisons (Figure 5) revealed that Arabs relied on right-sided visual areas more than Filipinos. Filipinos activated cerebellar regions more than Arabs during animal naming. There were no differences in activation between the groups during tool naming. Table 4 shows the LI values of activated regions of interest in Arab participants for animal and tool naming. Frontal processing was left-lateralized in most subjects during tool naming. Fewer subjects showed left lateralization during animal naming, which was bilateral in most subjects. The ANOVA test, performed to check for group-task interactions, showed a significant interaction with regards to the activation of bilateral OPs and temporo-occipital FFCs (Figure 6), with Arabs showing more prominent activations during tool naming compared to animal naming, and Filipinos eliciting more prominent activations during animal naming.

**Table (1):** Clusters and regions with significant activation during animal naming in comparison to the resting state in Arab and Filipino participants. The stated coordinates correspond with the area with maximum Z score (Z MAX) within each cluster. R: right; L: left; BL: bilateral; OPs: occipital poles; FFC: fusiform cortex; LG: lingual gyrus; PrC: precentral gyrus; IFG: inferior frontal gyrus; PO: pars opercularis; MFG: middle frontal gyrus; ACC: anterior cingulate cortex; SMA: supplementary motor area; OFC: orbitofrontal cortex; PT: pars triangularis; SFG: superior frontal gyrus; FP: frontal pole.

| Cluster Index | Voxels | $p$ | $-\log_{10}(p)$ | Z MAX | X (mm) | Y (mm) | Z (mm) | Regions |
| --- | --- | --- | --- | --- | --- | --- | --- | --- |
| **Arabs (Animal naming)** |  |  |  |  |  |  |  |  |
| 1 | 21612 | 0.000 | 99.7 | 7.08 | -14 | -98 | 10 | BL OPs, R FFC, BL LG |
| 2 | 232 | 0.000 | 3.62 | 4.18 | -50 | 2 | 52 | L PrC, L IFG (PO) |
| 3 | 149 | 0.004 | 2.36 | 4.09 | 42 | 6 | 28 | R PrC, R IFG (PO) |
| 4 | 115 | 0.016 | 1.78 | 4.39 | 44 | 2 | 62 | R MFG |
| 5 | 185 | 0.001 | 2.93 | 4.81 | -6 | 16 | 46 | BL ACC, L SMA |
| 6 | 118 | 0.015 | 1.84 | 4.48 | 34 | 20 | 4 | R insula |
| 7 | 103 | 0.027 | 1.57 | 5.56 | 20 | -28 | 0 | R thalamus |
| 8 | 156 | 0.003 | 2.48 | 4.02 | 30 | -68 | -52 | Cerebellum (R VIIb, VIIIa) |
| 9 | 308 | 0.000 | 4.64 | 5.01 | -36 | -62 | -54 | Cerebellum (L VIIb, VIIIa) |
| 10 | 193 | 0.001 | 3.05 | 4.71 | -26 | -36 | -42 | Cerebellum (L X, VI) |
| 11 | 190 | 0.001 | 3 | 5.65 | 22 | -36 | -42 | Cerebellum (R X) |
| **Filipinos (Animal naming)** |  |  |  |  |  |  |  |  |
| 1 | 21331 | 0.000 | 111 | 7.48 | 20 | -74 | -6 | BL LG, L FFC, L OP |
| 2 | 1839 | 0.000 | 25.6 | 5.15 | -46 | 4 | 38 | L PrC, L OFC, L MFG, L IFG (PT) |
| 3 | 503 | 0.000 | 9.89 | 4.64 | -4 | 22 | 44 | BL ACC, L SFG, L SMA |
| 4 | 275 | 0.000 | 6.11 | 4.66 | 34 | 26 | 4 | R IFG (PO), R insula |
| 5 | 256 | 0.000 | 5.76 | 4.85 | 42 | 8 | 32 | R PrC, R MFG |
| 6 | 182 | 0.000 | 4.28 | 4.52 | 46 | 36 | 18 | R FP |
| 7 | 97 | 0.005 | 2.29 | 4.49 | 32 | -8 | -8 | R putamen |
| 8 | 149 | 0.000 | 3.55 | 4.39 | -32 | -4 | -6 | L putamen |
| 9 | 287 | 0.000 | 6.32 | 5.86 | 22 | -28 | 2 | R thalamus |
| 10 | 107 | 0.003 | 2.55 | 5.93 | -18 | -30 | -2 | L thalamus |
| 11 | 150 | 0.000 | 3.58 | 4.62 | 20 | -34 | -42 | Cerebellum (R X) |
| 12 | 102 | 0.004 | 2.42 | 4.97 | -18 | -36 | -44 | Cerebellum (L X) |

**Table (2):** Clusters and regions with significant activation during tool naming in comparison to the resting state in Arab and Filipino participants. The stated coordinates correspond with the area with maximum Z score (Z MAX) within each cluster. R: right; L: left; BL: bilateral; LG: lingual gyrus; FFC: fusiform cortex; OPs: occipital poles; PrC: precentral gyrus; IFG: inferior frontal gyrus; MFG: middle frontal gyrus; PT: pars triangularis; FP: frontal pole; PO: pars opercularis; ACC: anterior cingulate cortex; SMA: supplementary motor area; SFG: superior frontal gyrus; ICC: intracalcarine cortex; LO: lateral occipital cortex; SPL: superior parietal lobule.

| Cluster Index | Voxels | p | -log<sub>10</sub>(p) | Z Max | x | y | z | Regions |
| --- | --- | --- | --- | --- | --- | --- | --- | --- |
| Arabs (Tool Naming) |  |  |  |  |  |  |  |  |
| 1 | 21331 | 0.000 | 111 | 7.48 | 20 | -74 | -6 | BL LG, BL FFC, R OP |
| 2 | 1038 | 0.000 | 13.6 | 4.92 | -50 | 0 | 50 | L PrC, L IFG (PO), L MFG |
| 3 | 235 | 0.000 | 4.19 | 4.63 | -48 | 34 | 14 | L IFG (PT), L FP |
| 4 | 147 | 0.002 | 2.7 | 4.62 | 40 | 10 | 32 | R MFG, R PrC, R IFG (PO) |
| 5 | 223 | 0.000 | 4 | 4.71 | -2 | 18 | 44 | L ACC, L SMA, L SFG |
| 6 | 150 | 0.002 | 2.75 | 5 | -30 | -64 | -50 | Cerebellum (L VIIb Left Crus II) |
| 7 | 119 | 0.007 | 2.16 | 5.66 | 22 | -38 | -44 | Cerebellum (R X) |
| 8 | 93 | 0.023 | 1.63 | 4.3 | 32 | -66 | -52 | Cerebellum (R VIIb R VIIIa) |
| 9 | 82 | 0.040 | 1.4 | 6.12 | -18 | -30 | 0 | L Thalamus |
| 10 | 79 | 0.050 | 1.33 | 5.7 | 20 | -28 | 0 | R Thalamus |
| Filipinos (Tool Naming) |  |  |  |  |  |  |  |  |
| 1 | 17748 | 0.000 | 113 | 6.94 | -2 | -94 | -4 | BL OPs, L LG, L ICC |
| 2 | 427 | 0.000 | 8.08 | 4.83 | 30 | -60 | 48 | BL LO, BL SPL |
| 3 | 1774 | 0.000 | 23.2 | 5.6 | -40 | 20 | 24 | L MFG, L IFG (PO and PT), L Insula, L PrC |
| 4 | 215 | 0.000 | 4.58 | 4.61 | 44 | 12 | 32 | R MFG, R IFG (PO), R PrC |
| 5 | 463 | 0.000 | 8.61 | 5.17 | -4 | 8 | 56 | L SMA, L SFG, BL ACC |
| 6 | 148 | 0.000 | 3.25 | 4.66 | 32 | 24 | 8 | R IFG (PO) |
| 7 | 80 | 0.022 | 1.67 | 3.77 | 10 | -16 | 4 | R Thalamus |
| 8 | 101 | 0.007 | 2.19 | 4.73 | -14 | 0 | 14 | L Caudate, L Putamen |
| 9 | 70 | 0.040 | 1.4 | 4.05 | -10 | -4 | 10 | L Thalamus, L Putamen, L Thalamus |

**Figure (2):**
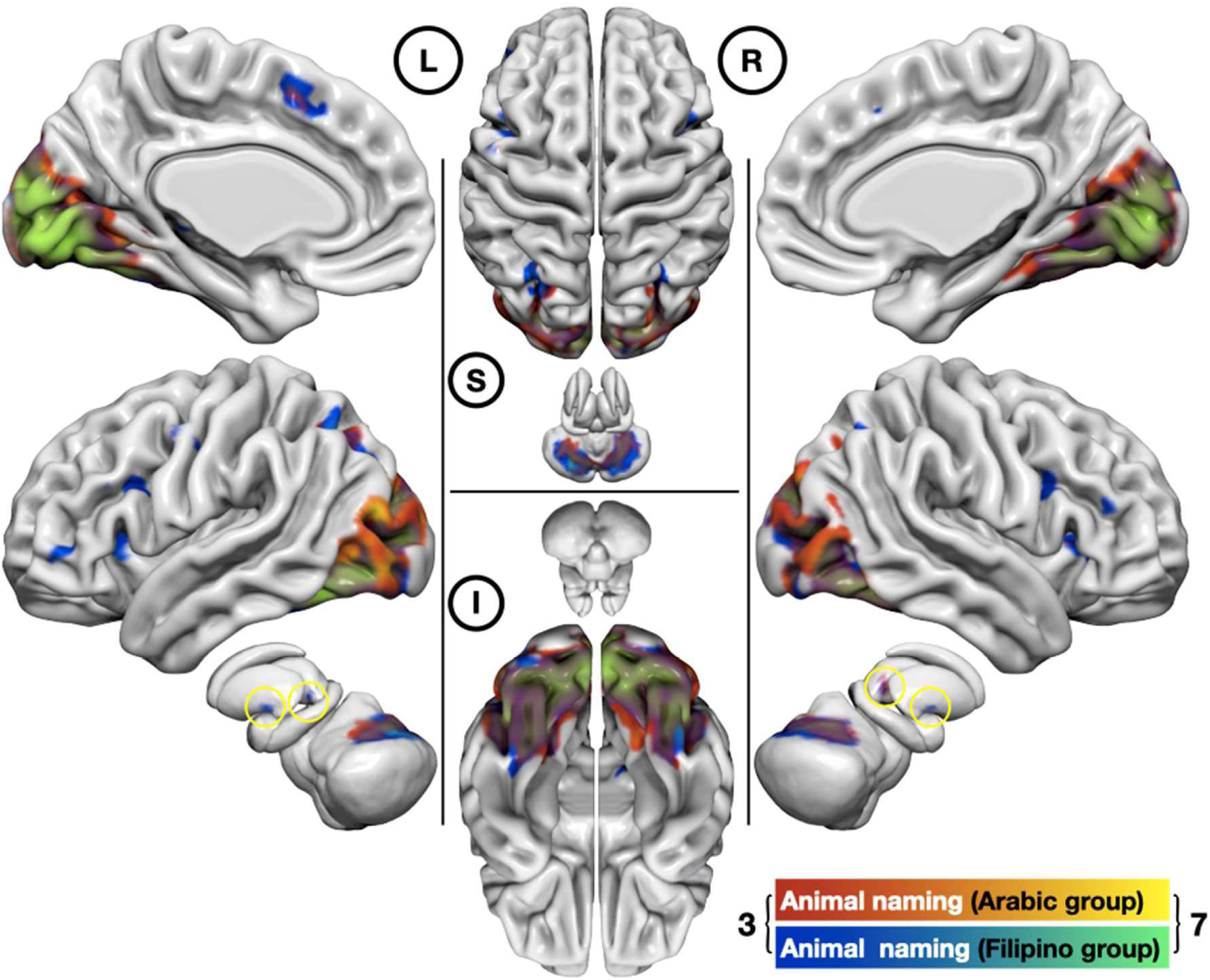
Group average maps of activations during the animal naming task for both Arabs and Filipinos (red-yellow and blue-green color bars, respectively) superimposed on brain surface meshes. The color bars show fMRI signal levels (Z-scores) above the 0.05 significance threshold. L, R, S, and I are the left, right, superior, and inferior views respectively.

**Figure (3):**
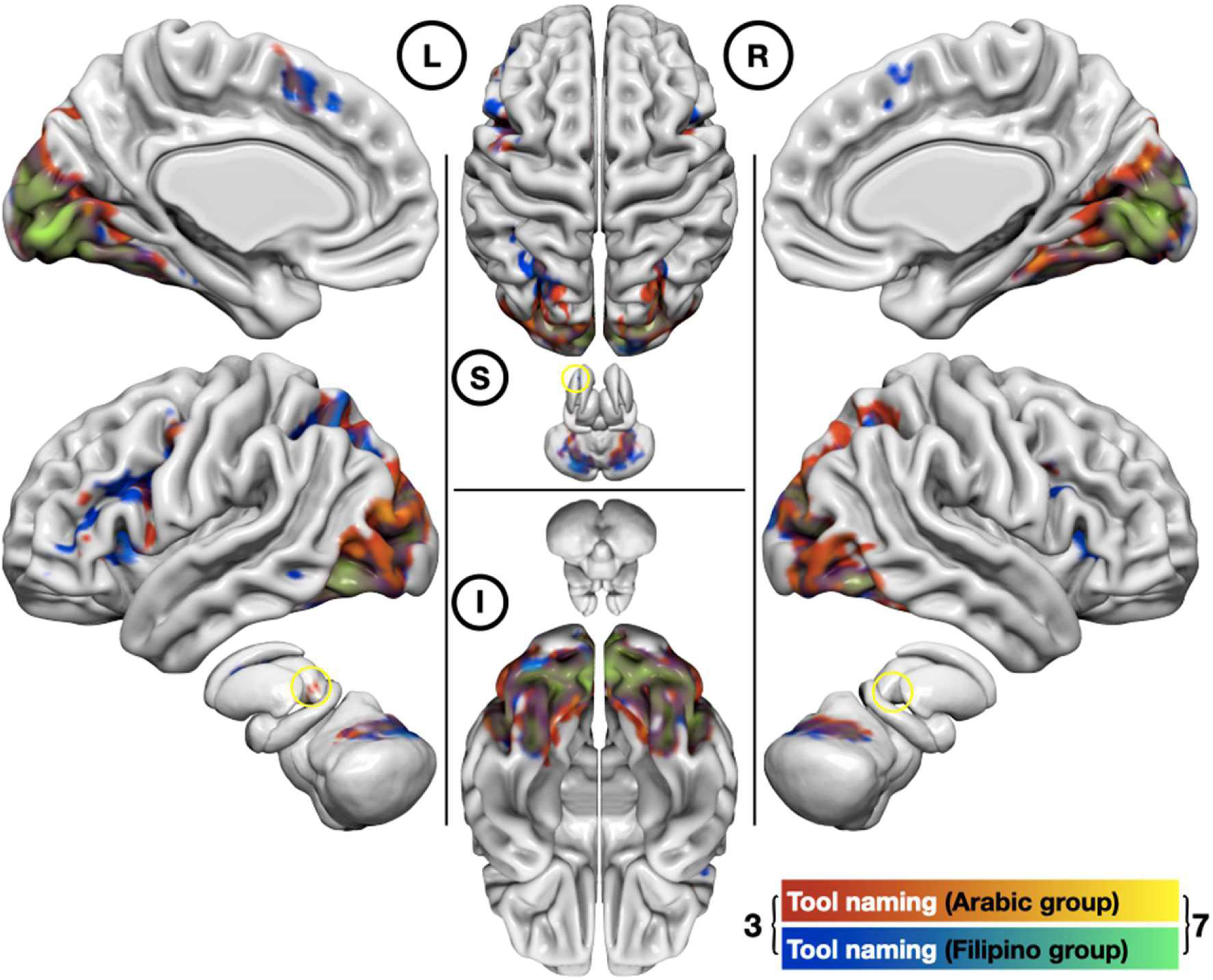
Group average maps of the activations during tool naming for Arabs and Filipinos (red-yellow and blue-green color bars, respectively), superimposed on brain surface meshes. The color bars show fMRI signal levels (Z-scores) above the 0.05 significance threshold. L, R, S, and I are the left, the right, the superior, and inferior views, respectively.

**Table (3):** Clusters and regions shown with significant activation during cross-task and cross-group comparisons. The stated coordinates correspond with the area with maximum Z score within each cluster (Z MAX). R: right; L: left; BL: bilateral; LG: lingual gyrus; FFC: fusiform cortex; OPs: occipital poles; PrC: precentral gyrus; IFG: inferior frontal gyrus; MFG: middle frontal gyrus; PT: pars triangularis; FP: frontal pole; PO: pars opercularis; ACC: anterior cingulate cortex; SMA: supplementary motor area; SFG: superior frontal gyrus; ICC: intracalcarine cortex; LO: lateral occipital cortex; SPL: superior parietal lobule. * The condition (tool > animal naming) yielded no significant results in Filipinos. ** The condition (Filipinos > Arabs) showed no significant difference for tool naming. *** There is no significant run effect.

| Cluster Index | Voxels | *P* | $-\log_{10}(P)$ | Z MAX | Z-MAX X (mm) | Z-MAX Y (mm) | Z-MAX Z (mm) | Region Name |
| --- | --- | --- | --- | --- | --- | --- | --- | --- |
| **Arabs (Animal > Tool naming)** |  |  |  |  |  |  |  |  |
| 1 | 1793 | 0.000 | 20.2 | 5.37 | 4 | -92 | 18 | BL OPs |
| 2 | 413 | 0.000 | 6.75 | 5.38 | 52 | -72 | 4 | R LO (inferior division) |
| 3 | 88 | 0.030 | 1.53 | 4.98 | 40 | -74 | -12 | R FFC (temporo-occipital) |
| **Arabs (Tool > Animal naming)** |  |  |  |  |  |  |  |  |
| 1 | 132 | 0.004 | 2.42 | 4.02 | -22 | -52 | 62 | L SPL |
| 2 | 118 | 0.007 | 2.15 | 4.19 | 22 | -50 | 70 | R SPL |
| 3 | 84 | 0.040 | 1.44 | 4.2 | -38 | -36 | 44 | L postcentral gyrus |
| **Filipinos (Animal > Tool naming)** |  |  |  |  |  |  |  |  |
| 1 | 2730 | 0.000 | 35.5 | 5.32 | 4 | -84 | 16 | R ICC, R OP |
| 2 | 296 | 0.000 | 6.92 | 5.08 | -46 | -78 | 0 | L LO (inferior division), L FFC |
| 3 | 82 | 0.009 | 2.04 | 4.44 | 42 | -44 | -18 | R FFC (temporo-occipital) |
| **Filipinos (Tool > Animal)\*** |  |  |  |  |  |  |  |  |
| 1 | -- | < 0.05 | -- | -- | -- | -- | -- | Not significant at $p < 0.05$ level for ROIs |
| **Animal naming (Arabs > Filipinos)** |  |  |  |  |  |  |  |  |
| 1 | 110 | 0.019 | 1.72 | 3.95 | 22 | -50 | -10 | R LG, R FFC (occipital), R LG |
| **Animal naming (Filipinos > Arabs)** |  |  |  |  |  |  |  |  |
| 1 | 268 | 0.000 | 4.3 | 4.66 | 24 | -72 | -26 | Cerebellum (R Crus) |
| 2 | 114 | 0.016 | 1.79 | 4.04 | -32 | -72 | -30 | Cerebellum (L Crus) |
| **Tool naming (Arabs > Filipinos) \*\*** |  |  |  |  |  |  |  |  |
| 1 | 112 | 0.011 | 1.95 | 4.26 | 28 | -70 | -6 | R FFC |
| **Group-Run interaction (Arabic vs Filipino) x (tool vs animal)** |  |  |  |  |  |  |  |  |
| 1 | 4250 | 0.000 | 37.7 | 5.87 | 10 | -92 | -2 | R OP |
| 2 | 140 | 0.002 | 2.65 | 4.9 | 42 | -44 | -18 | R FFC (temporo-occipital) |
| 3 | 105 | 0.011 | 1.95 | 4.41 | -44 | -48 | -22 | L FFC (temporo-occipital) |
| 4 | 80 | 0.039 | 1.41 | 4.58 | -46 | -78 | 0 | L LO (inferior division) |

\* In Filipinos the Tool naming > Animal naming condition showed no significant results.
\*\* In tool naming, the Filipinos > Arabs condition showed no significant results.

**Figure (4):**
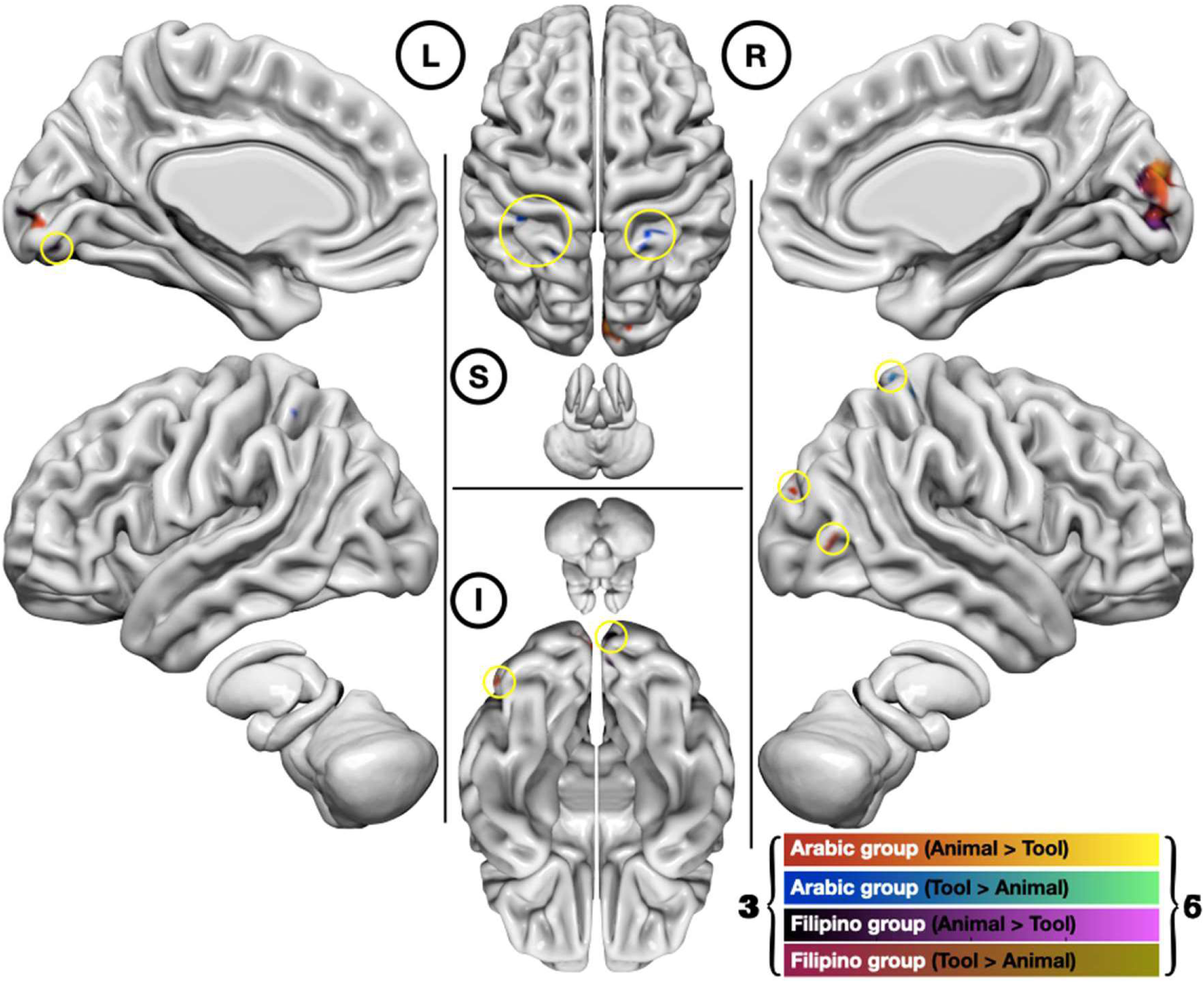
Paired t-test maps of animal and tool naming tasks with cross task comparisons for Arab and Filipino participants (color coding as indicated), superimposed on brain and cerebellum surface meshes. The color bars show fMRI signal level (Z-scores) above the 0.05 significance threshold. L, R, S, and I are the left, the right, the superior, and inferior views, respectively.

**Figure (5):**
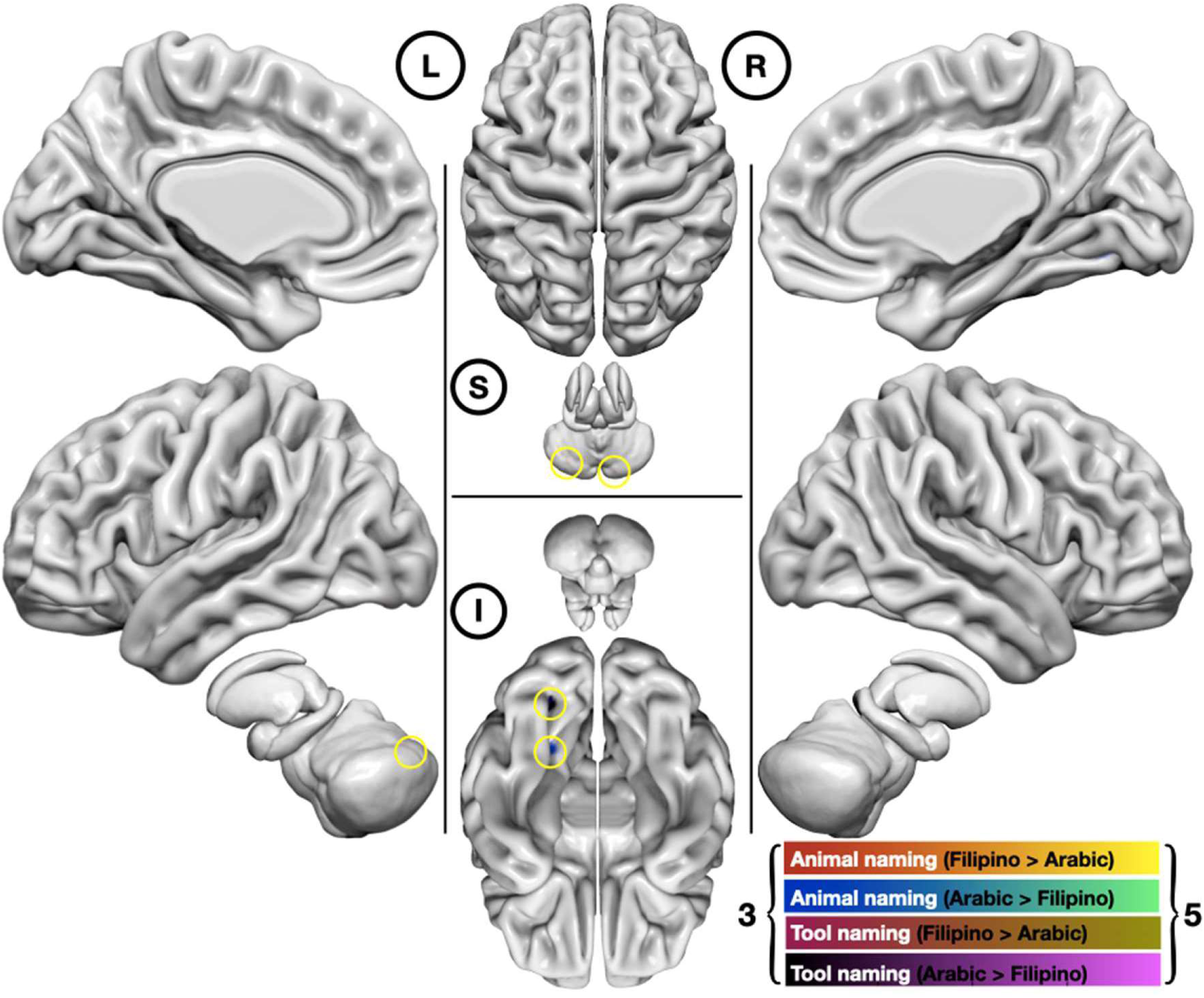
Paired t-test maps of cross-group comparisons between Arab and Filipino participants during each of the tasks (color coding as indicated), superimposed on brain and cerebellum surface meshes. L, R, S, and I are the left, the right, the superior, and inferior views respectively). The color bars show fMRI signal level (Z-scores) above the 0.05 significance threshold.

**Table 4:** Laterality Indices of activations in cortical areas of interest in 18 Arab subjects. ICC: intracalcarine cortex; LO: lateral occipital cortex; FFC: fusiform cortex; OP: occipital pole; FP: frontal pole; IFG (OP): inferior frontal gyrus, pars opercularis; IFG (TG): inferior frontal gyrus, pars triangularis); SFG: superior frontal gyrus; MFG: middle frontal gyrus; ACC: anterior cingulate cortex; PrCG: precentral gyrus; SMA: supplemental motor area. * left lateralization; ** right lateralization.

|  | ICC | LO | Lingual | FFC | OP | FP | IFG (PO) | IFG (PT) | SFG | MFG | ACC | PrCG | SMA |
| --- | --- | --- | --- | --- | --- | --- | --- | --- | --- | --- | --- | --- | --- |
| **Animal Naming** |  |  |  |  |  |  |  |  |  |  |  |  |  |
| 1 | 0.05 | -0.02 | -0.03 | 0.04 | 0.01 | 0.10 | 0.00 | 0.05 | -0.08 | 0.05 | 0.18 | 0.02 | -0.24** |
| 2 | -0.04 | 0.04 | 0.03 | 0.24* | 0.01 | -0.22** | -0.04 | -0.49** | 0.13 | -0.04 | 0.03 | 0.04 | 0.40* |
| 3 | -0.23** | 0.01 | -0.08 | 0.06 | 0.06 | -0.09 | 0.28* | -0.17 | -0.09 | -0.19 | -0.06 | 0.05 | -0.89 |
| 4 | -0.04 | 0.07 | 0.02 | 0.01 | 0.02 | -0.12 | 0.67* | 0.50* | 0.48* | 0.47* | 0.37* | 0.41* | 0.39* |
| 5 | -0.06 | 0.11 | 0.02 | 0.14 | 0.03 | 0.36* | 0.55* | 0.48* | 0.53* | 0.49* | 0.95 | 0.29* | 0.14 |
| 6 | 0.10 | 0.03 | 0.01 | 0.24* | -0.04 | -0.12 | -0.09 | -0.05 | -0.05 | -0.13 | 0.51* | -0.02 | 0.12 |
| 7 | -0.05 | 0.04 | -0.05 | 0.13 | 0.05 | 0.21* | 0.33* | 0.18 | 0.35* | 0.19 | 0.48* | 0.16 | -0.11 |
| 8 | 0.00 | 0.03 | -0.02 | 0.01 | 0.06 | 0.09 | 0.19 | 0.26* | 0.26* | 0.26* | 0.21* | 0.23* | 0.13 |
| 9 | -0.13 | 0.00 | -0.06 | 0.11 | -0.02 | -0.25** | -0.34** | -0.29** | -0.17 | -0.30** | -0.13 | -0.39** | -0.15 |
| 10 | -0.02 | 0.06 | -0.04 | 0.09 | 0.08 | -0.56** | -0.28** | -0.56** | 0.26* | -0.32** | -0.08 | -0.17 | -0.62** |
| 11 | -0.01 | 0.04 | -0.01 | 0.10 | 0.08 | 0.11 | -0.09 | -0.13 | 0.14 | 0.05 | 0.29* | -0.12 | 0.12 |
| 12 | -0.05 | 0.06 | 0.03 | 0.11 | 0.08 | -0.02 | 0.21* | 0.15 | 0.03 | 0.18 | -0.11 | 0.06 | -0.02 |
| 13 | -0.01 | 0.04 | 0.01 | 0.32* | 0.02 | -0.15** | -0.08 | -0.19 | 0.05 | -0.13 | -0.17 | 0.03 | 0.37* |
| 14 | -0.03 | 0.02 | -0.03 | -0.05 | -0.06 | -0.57* | -0.66** | -0.78 | -0.87 | -0.77 | -0.48* | -0.90 | -0.33** |
| 15 | -0.05 | 0.07 | -0.04 | 0.36* | 0.05 | 0.01 | -0.02 | -0.14 | -0.16 | -0.15 | -0.19 | -0.11 | -0.25** |
| 16 | -0.04 | -0.02 | -0.02 | 0.53* | 0.03 | 0.34* | 0.31* | 0.34* | 0.51* | 0.36* | 0.45* | 0.31* | 0.78 |
| 17 | -0.07 | 0.06 | -0.01 | 0.14 | 0.05 | -0.30** | -0.02 | -0.12 | -0.25* | -0.20 | 0.00 | -0.07 | 0.35* |
| 18 | 0.01 | -0.02 | -0.01 | 0.27* | -0.08 | -0.03 | 0.33* | 0.31* | 0.33* | 0.24* | 0.35* | 0.07 | 0.39* |
| **Tool Naming** |  |  |  |  |  |  |  |  |  |  |  |  |  |
| 1 | 0.11 | 0.01 | -0.01 | 0.04 | -0.01 | 0.00 | -0.10 | -0.05 | 0.11 | 0.19 | 0.20* | 0.03 | -0.30** |
| 2 | -0.03 | 0.09 | 0.01 | 0.20 | 0.02 | 0.48* | 0.39* | 0.46* | 0.44* | 0.38* | 0.38* | 0.38* | 0.77* |
| 3 | -0.20** | 0.14 | -0.10 | 0.22* | 0.10 | -0.42* | 0.38* | -0.26** | 0.33* | 0.25* | 0.26* | 0.27* | -0.74** |
| 4 | -0.07 | 0.17 | -0.04 | 0.17 | 0.05 | 0.34* | 0.54* | 0.54* | 0.38* | 0.52* | 0.18 | 0.39* | 0.70* |
| 5 | -0.05 | 0.16 | 0.09 | 0.92* | 0.04 | 0.06 | 0.55* | 0.46* | -0.03 | 0.33* | 0.05 | 0.23* | 0.31* |
| 6 | 0.09 | 0.05 | -0.04 | 0.27* | -0.03 | 0.13 | 0.40* | 0.38* | 0.19 | 0.24* | 0.33* | 0.25* | 0.52* |
| 7 | -0.07 | 0.12 | -0.13 | 0.20* | 0.06 | 0.38* | 0.63* | 0.53* | 0.50* | 0.42* | 0.89* | 0.27* | 0.73* |
| 8 | 0.00 | 0.07 | -0.03 | 0.15 | 0.06 | 0.58* | 0.59* | 0.59* | 0.49* | 0.61* | 0.33* | 0.37* | 0.55* |
| 9 | -0.13 | -0.01 | -0.05 | 0.21* | -0.01 | 0.24* | 0.25* | 0.28* | 0.05 | 0.16 | 0.11 | 0.06 | 0.34* |
| 10 | -0.02 | 0.05 | -0.01 | 0.24* | 0.10 | 0.05 | 0.31* | 0.26* | 0.20 | 0.19 | 0.12 | 0.25* | 0.76* |
| 11 | -0.04 | 0.07 | -0.04 | 0.03 | 0.02 | 0.29* | 0.14 | 0.17 | 0.28* | 0.24* | 0.26* | -0.02 | 0.07 |
| 12 | -0.10 | -0.02 | 0.00 | 0.17 | -0.01 | 0.32* | 0.53* | 0.56* | 0.12 | 0.35* | 0.48* | -0.14 | 0.20 |
| 13 | -0.01 | 0.10 | 0.02 | 0.36* | 0.04 | -0.13 | 0.39* | 0.01 | 0.29* | 0.05 | 0.26* | 0.35* | 0.76* |
| 14 | -0.06 | 0.11 | -0.02 | -0.12 | 0.02 | -0.33* | -0.53** | -0.42** | -0.83** | -0.50** | -0.85** | -0.61** | -0.33** |
| 15 | -0.05 | 0.04 | -0.02 | 0.25* | 0.04 | -0.47* | -0.68** | -0.64** | -0.57** | -0.70** | -0.24** | -0.50** | -0.83** |
| 16 | -0.07 | 0.12 | -0.01 | 0.58* | 0.07 | 0.41* | 0.34* | 0.44* | 0.41* | 0.31* | 0.56* | 0.23* | 0.43* |
| 17 | -0.14 | 0.03 | -0.06 | 0.15 | 0.04 | -0.03 | 0.07 | 0.10 | 0.09 | 0.06 | 0.13 | -0.06 | 0.20* |
| 18 | 0.12 | 0.14 | 0.10 | 0.37* | -0.02 | 0.08 | 0.76* | 0.92* | 0.87* | 0.87* | 0.94* | 0.49* | 0.91* |

**Figure (6):**
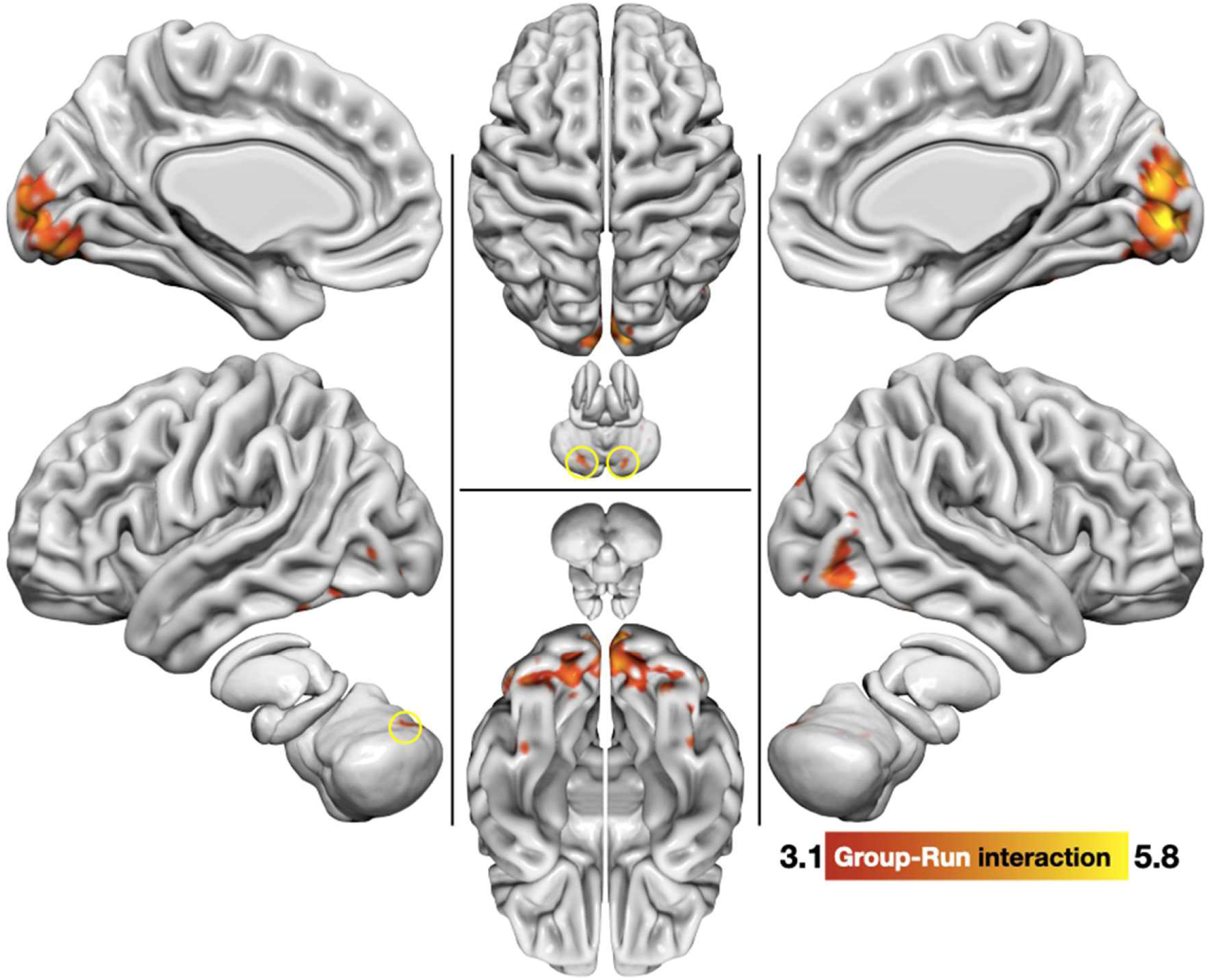
Group-task interaction map resulted from repeated measures ANOVA is superimposed on brain and cerebellum surface meshes, where L, R, S, and I are the left, the right, the superior, and inferior views respectively). The color bars show fMRI signal level (Z-scores) above the 0.05 significance threshold.

## DISCUSSION

This study demonstrated the activation of primary and secondary visual association cortices in association with picture-naming by Arab and Filipino participants. Although both groups activated these visual regions bilaterally, Arab participants recruited right-sided visual areas (the LO and FFC) more strongly than the Filipino group. This reliance on right-sided visual areas by Arab participants was particularly evident during animal naming as shown in the cross-task comparison. It suggests that Arabs have a culture-specific pattern of processing pictures of animals in the right ventral visual stream and may relate to the previously described role of the right visual system in processing pictures with faces (Rossion et al., 2003). It may be that Arabs require recruitment of visual areas more so than Filipinos, due to lack of direct experience with the various species of animals presented to participants, which may be supported by the difference in lexical frequency that these words are presented in corpora for each of the languages that may underlie divergent psycholinguistic processing by the two groups. The Philippines is a largely forested, and tropical region, lending access to hundreds of species of mammals and a diverse fauna (Oliver, 1994), whereas, Saudi Arabia is a typical desert climate that has witnessed a decline in animal diversity from the mid-nineteenth to mid-twentieth century (Buttiker et al., 1990). In the same vein, Alanazi (2018) showed that Saudi children generally do not have adequate forms of reasoning for biological classification, as “half of the Saudi children in the study thought that bats, bees, and butterflies were birds”. These findings highlight that children in Saudi primary schools show a significant range of alternative conceptions regarding the categorization of animals (Alanazi, 2018).

Moreover, the Philippines has 200 hundred zoos where Filipinos may be exposed to the exotic animals presented in the picture naming task (Almazan et al., 2005), compared to approximately seven zoos in Saudi Arabia (Mohammed, 2013). Filipinos appeared to recruit cerebellar regions more than Arabs during animal naming which may relate to the cerebellum’s role in effective word retrieval, moreover, as the Filipino cohort is entirely bilingual (Tagalog-English), the cerebellum’s activity may be related to verbal interference found in bilinguals (Filippi et al., 2020; Gollan & Acenas, 2004; Mariën et al., 2014). During tool naming and in contrast to animal naming, Arabs activated a predominantly left-lateralized network. This included areas known to sub-serve praxis and relate to retrieval of kinesthetic properties of tools that are represented along the dorsal visual stream (Moll et al., 2000; Shimotake et al., 2017).

Temporo-limbic (mesial temporal, entorhinal and peri-rhinal, amygdala) activation during animals, a previously reported finding in studies with Western participants, was notably absent in our study. This may be related to the nature of the naming task used in our study, the pictures may have been too simple and the task too low in complexity to engage participants enough to elicit a subcortical response. An alternate possibility is that this is a culturally-dependent phenomenon. The animals shown may not have carried the same degree of emotional salience to our participants as they did to Western subjects in prior studies, although this remains a speculation without a direct comparative study of Western and Arab subjects doing this task. Gutchess and colleagues (2006) provided an example of a culture-dependent neural response to picture presentations. In comparing Chinese and American participants, it was revealed that Chinese participants activated a fronto-parietal network implicated in executive control, whereas American subjects activated temporal and cingulate areas thought to relate higher order visual processing of details. This was attributed to the emphasis of Asian cultures on context and relationships of objects, which contrasts with American culture’s focus on individual entities (Gutchess et al., 2006). Our findings may indicate that picture naming of animate and inanimate objects depends on specific cultural-lingual properties, and would caution against generalizing findings of the more commonly studied populations on participants from other cultures. This is especially relevant in high-risk procedures, such as neurosurgery (Chakraborty & McEvoy, 2008).

## LIMITATIONS

The task was thoroughly explained to the participants to ensure they performed covert (silent) naming correctly, however, participants were not asked to practice the task outside the scanner to ensure accurate performance of the task. This study employed covert naming, which is not designed to detect naming errors, as such, brain activation maps do not distinguish error related activity. While the variance in lexical frequencies of the picture names between the two languages may act as a potential cofound to the results. In this study a block design was used but timestamps for each stimulus picture was not recorded, and thus it was not possible to include the lexical frequencies as covariates in the analysis. Even though the stimuli were selected from an extensively studied picture bank in research using picture naming tasks, these psycholinguistic differences may yield significant findings in future work. Furthermore, no measures of naming latency or naming accuracy were obtained, such factors may be impacted by the lexical frequency of the words and may be further confounded by bilingualism. Future studies may control for these properties, and may even seek examine differences between stimuli with matched and non-matched psycholinguistic properties

## CONCLUSIONS

This study is the first study to demonstrate fMRI correlates of picture naming in Arab individuals, compared to Filipino participants, in categories of animal and tool naming. The findings illustrate category specificity of neural activation to the semantic content of the pictures. Whereas both groups activated primary and secondary visual areas, animal naming tended to activate nondominant-hemisphere ventral visual areas in Arabs, and bilateral visual and cerebellar regions in Filipinos. In Arabs, tool pictures activated predominantly dominant-hemisphere fronto-parietal areas fitting with the sensorimotor and kinesthetic properties of tools, whereas tool naming was not associated specific activity in Filipino participants, and no differences between groups were founds. Our study did not demonstrate significant temporo-limbic activation with animal naming, a finding that may require further study to illustrate whether it is culture-dependent or related to the task used in this study. This study used a simple covert naming task, whereas tests that are high in complexity, and task demand may differ in results.

