## Supplementary Table S1 for "Category-specific fMRI correlates of picture naming: A study with Arabs and Filipinos"

Supplementary Materials.

Table S1. Names of animal pictures in Arabic and Tagalog with corresponding lexical frequencies per one million.

| Animal Picture | Arabic Frequency | Animal Name in Arabic | Tagalog Frequency | Animal Name in Tagalog |
|---|---|---|---|---|
| Alligator | 0,16 | تِمسَاح | 12,30 | Buwaya |
| Bear | 2,16 | دب | 3,52 | Oso |
| Bird | 1,46 | طير | 49,64 | Ibon |
| Camel | 2,94 | جمل | 29,15 | Kamelyo |
| Cat | 0,96 | قطة | 6,87 | Cat |
| Chicken | 0,75 | دجاجة | 100,59 | Manok |
| Cow | 1,51 | بقرة | 174,26 | Baka |
| Deer | 1,64 | غزال | 14,89 | Deer |
| Dog | 2,94 | كلب | 90,48 | Aso |
| Donkey | 1,69 | حمار | 4,25 | Asno |
| Duck | 0,49 | بَطَّهْ | 36,54 | Pato |
| Eagle | 1,25 | نِسر | 3,71 | Eagle |
| Elephant | 2,42 | فِيلْ | 0,45 | Elephant |
| Fish | 2,03 | سمكة | 60,07 | Isda |
| Fox | 2,21 | ثعلب | 0,32 | Soro |
| Frog | 0,13 | ضفدع | 10,16 | Palaka |
| Giraffe | 0,1 | زرافهْ | 0,25 | Giraffe |
| Goat | 0,08 | مَعْزَهْ | 11,33 | Kambing |
| Gorilla | 0,03 | غُورِلَّا | 2,24 | Bakulaw |
| Horse | 4,58 | حصَانْ | 27,91 | Kabayo |
| Kangaroo | NA | كُنْغُر | 0,68 | Kanggaroo |
| Leopard | 64,22 | فهد | 0,52 | Leopardo |
| Lion | 0,6 | اسد | 41,09 | Leon |
| Monkey | 0,47 | قِرْد | 1,67 | Matsing |
| Mouse | 1,38 | فَارْ | 1,34 | Mouse |
| Ostrich | 0,39 | نعَامَهْ | NA | Ostrich |
| Owl | 0,1 | بُومَهْ | 1,14 | Kuwago |
| Peacock | 0,16 | طاووس | 0,17 | Peacock |
| Penguin | 8,95 | بَطْرِيقْ | 0,86 | Penguin |
| Pig | 0,42 | خِنْزِيرْ | 55,56 | Baboy |
| Rabbit | 0,05 | أَرْنِبْ | 4,52 | Kuneho |
| Racoon | NA | راكون | 0,04 | Racoon |
| Rhinoceros | 16,13 | وَحِيدِ القَرْن | NA | Rhinoceros |
| Rooster | 16,13 | ديك | 0,54 | Rooster |
| Sheep | 0,68 | خروف | 23,86 | Tupa |
| Snake | 0,81 | ثعبان | 26,25 | Ahas |

| | | | | |
|---|---|---|---|---|
| Squirrel | 0,26 | سِنْجَابْ | 0,17 | ardilya |
| Tiger | 12,43 | نِمر | 3,84 | Tigre |
| Turtle | 0,39 | سلحفاة | NA | Pagong |
| Zebra | 1,69 | حِمَارْ وحْشِي | 0,11 | Zebra |

| Animal | | |
|---|---|---|
| | Arabic | Tagalog |
| *M* | 4,07 | 21,66 |
| *SD* | 10,81 | 36,18 |
| *Range* | 64,19 | 174,22 |
| *Min* | 0,03 | 0,035 |
| *Max* | 64,22 | 174,26 |

The names and lexical frequencies for Arabic are based on the Aralex corpus, the names presented are in Modern Standard Arabic (MSA) form, and the frequencies are reported per million. The corpus contains 40,000,000 words. The names and lexical frequencies for Tagalog are based on the http://www.SketchEngine.eu website (Tagalog (Filipino) Web 2019 (tlTenTen19) corpus, the frequencies presented are per million. The corpus contains 198,303,250 words.

Table S2. Names of tool pictures in Arabic and Tagalog with corresponding lexical frequencies per one million.

| Tool Picture | Arabic Frequency | Tool Name in Arabic | Tagalog Frequency | Tool Name in Tagalog |
|---|---|---|---|---|
| ashtray | 0,03 | مرمدة | 0,43 | ashtray |
| axe | 0,21 | فَأسْ | 2,19 | palakol |
| basket | 8,50 | سلّة | 2,17 | basket |
| bell | 3,49 | جرس | 1,34 | kampanilya |
| bottle | 2,89 | زجاجة | 39,26 | bote |
| broom | 0,29 | مكنسة | 4,42 | walis |
| brush | 0,81 | فرشاة | NA | magsipilyo |
| candle | 2,52 | شمعةْ | 14,81 | kandila |
| chain | 78,60 | سِلْسَلَةْ | 8,12 | kadena |
| chair | 8,22 | كرسي | 67,36 | upuan |
| comb | 0,21 | مشط | 0,06 | magsuklay |
| cup | 2,96 | فِنْجانْ | 80,81 | tasa |
| fork | 2,68 | شوكة | NA | tinidor |
| frying pan | 0,05 | مقلاة | 10,27 | kawali |
| glove | 0,31 | قفاز | 2,40 | guwantes |
| hammer | 1,33 | مطَرْقَة | 3,57 | martilyo |
| hanger | 1,22 | شماعة | 0,06 | hanger |
| iron | 15,68 | حديدْ | 28,83 | bakal |
| key | 13,76 | مِفْتاح | 42,55 | susi |
| knife | 1,64 | سكين | 12,06 | kutsilyo |
| ladder | 29,05 | سلم | 19,19 | hagdan |
| lock | 0,81 | قفل | 1,22 | lock |
| nail | 1,09 | مسمار | 18,10 | kuko |
| needle | NA | إبْرهْ | 8,03 | karayom |
| paintbrush | 0,81 | فرشاة الرسم | 9,53 | pintura |
| pen | 5,33 | قلم | 11,45 | panulat |
| pencil | NA | قلَمْ رصَاصْ | 10,71 | lapis |
| pliers | 0,47 | كماشة | NA | mga plier |
| plug | 0,86 | قابس | 0,42 | plug |
| pot | 3,88 | وعاء | 7,62 | palayok |
| ruler | 0,39 | مسطرة | 83,64 | pinuno |
| saw | 0,03 | مِنْشَارْ | 2,22 | nakita |
| scissors | 0,36 | مقَصٌ | 4,83 | gunting |
| screwdriver | 0,03 | مفك | 0,05 | distornilyador |
| spoon | 0,52 | ملعقة | 14,95 | kutsara |
| stool | NA | كرسي مرتفع | NA | dumi ng tao |

| | | | | |
|---|---|---|---|---|
| telephone | 6,14 | هاتف | 71,32 | telepono |
| watch | NA | ساعة يد | 0,54 | manood |
| whistle | 1,85 | صفارة | 0,85 | sipol |
| wrench | 13,76 | مِفْتاحْ | 0,14 | wrench |

Tool Names

| | Arabic | Tagalog |
|---|---|---|
| *M* | 5,86 | 16,26 |
| *SD* | 13,8258589 | 23,7882565 |
| *Range* | 78,57 | 83,59 |
| *Min* | 0,03 | 0,05 |
| *Max* | 78,60 | 83,64 |

The names and lexical frequencies for Arabic are based on the Aralex corpus, the names presented are in Modern Standard Arabic (MSA) form, and the frequencies are reported per million. The corpus contains 40,000,000 words. The names and lexical frequencies for Tagalog are based on the http://www.SketchEngine.eu website (Tagalog (Filipino) Web 2019 (tlTenTen19) corpus, the frequencies presented are per million. The corpus contains 198,303,250 words.
